# Lymphocyte activation gene 3 (Lag3) supports Foxp3^+^ Treg cell function by restraining c-Myc-dependent aerobic glycolysis

**DOI:** 10.1101/2023.02.13.528371

**Authors:** Dongkyun Kim, Giha Kim, Rongzhen Yu, Juyeun Lee, Sohee Kim, Kevin Qiu, Elena Montauti, Deyu Fang, Navdeep S. Chandel, Jaehyuk Choi, Booki Min

## Abstract

Lymphocyte activation gene 3 (Lag3) has emerged as the next-generation immune checkpoint molecule due to its ability to inhibit effector T cell activity. Foxp3^+^ regulatory T (Treg) cells, a master regulator of immunity and tolerance, also highly express Lag3. While Lag3 is thought to be necessary for Treg cell-mediated regulation of immunity, the precise roles and underlying mechanisms remain largely elusive. In this study, we report that Lag3 is indispensable for Treg cells to control autoimmune inflammation. Utilizing a newly generated Treg cell specific Lag3 mutant mouse model, we found that these animals are highly susceptible to autoimmune diseases, suggesting defective Treg cell function. Genome wide transcriptome analysis further uncovered that Lag3 mutant Treg cells upregulated genes involved in metabolic processes. Mechanistically, we found that Lag3 limits Treg cell expression of Myc, a key regulator of aerobic glycolysis. We further found that Lag3-dependent Myc expression determines Treg cells’ metabolic programming as well as the in vivo function to suppress autoimmune inflammation. Taken together, our results uncovered a novel function of Lag3 in supporting Treg cell suppressive function by regulating Myc-dependent metabolic programming.

## Introduction

Foxp3^+^ regulatory T (Treg) cells are indispensable regulators of immunity (Sakaguchi et al., 2008). Due to their potent suppressive properties, their function must be tightly controlled. Diminished Treg cell function could result in uncontrolled immune activation seen in chronic inflammation or in autoimmunity, while enforced Treg cell function may interfere with immune responses as seen in intratumoral Treg cells’ inhibition of antitumor effector immunity (Vignali et al., 2008). Treg cells express a battery of surface receptors capable of modulating Treg cell functions. For example, ICOS supports Treg cells’ proliferation, survival, and function partly through IL-10 induction (Busse et al., 2012; Kornete et al., 2012; Lohning et al., 2003), although ICOS is also shown to limit Treg cell accumulation and function especially within visceral adipose tissue (Mittelsteadt et al., 2021). PD-1 plays a role to enhance Foxp3 expression and their suppressive activity (Gotot et al., 2012; Lin et al., 2019), while PD-1-deficient Treg cells may elicit more potent suppressive activity and ameliorate autoimmune inflammation (Tan et al., 2021).

Lymphocyte activation gene 3 (Lag3) is a CD4-like coreceptor expressed on activated T cells, including Treg cells (Andrews et al., 2017). Analogous to CTLA-4, Lag3 reportedly limits conventional T cell proliferation and IL-2 production, acting as an inhibitory coreceptor (Burnell et al., 2022). As for the role of Lag3 in Treg cells, earlier study reported that Lag3 is necessary for optimal Treg cell suppressive function (Huang et al., 2004). Lag3^+^ Treg cells found in the tumor tissues display highly activated phenotypes and produce immunosuppressive cytokines such as IL-10 and TGFβ (Camisaschi et al., 2010; Gagliani et al., 2013). We also found that Lag3-deficient Treg cells are unable to mitigate autoimmune inflammation following transfer into mice with ongoing experimental autoimmune encephalomyelitis (EAE) (Kim et al., 2019). However, the precise mechanisms underlying Lag3-dependent regulation of Treg cell function remain largely elusive.

Increasing evidence suggests that distinct metabolic profiles are wired into different immune cell subsets and their functions (Ganeshan and Chawla, 2014). Treg cells utilize metabolic programs distinct from those used by effector lineage cells (Shi and Chi, 2019). Similar to naive T cells, Treg cells primarily exploit oxidative phosphorylation as the major energy source (Berod et al., 2014). Enhanced aerobic glycolysis, through which pyruvate is converted to lactate rather than enters the TCA cycle, is largely used by cancer cells or by effector T cells and is shown to inhibit Treg cell function (Shyer et al., 2020). Myc, an oncogene directly regulating genes involved in glycolytic activity, is tightly controlled in Treg cells (Dong et al., 2020; Min et al., 2022), and Foxp3, the master transcription factor for Treg cells, is critically involved in reprograming Treg cell metabolic profiles in part by suppressing Myc expression (Angelin et al., 2017).

In the current study, we investigated the roles of Lag3 on Treg cells by using a newly generated Treg cell-specific Lag3-deficient animal model and found that Lag3 expression in Treg cells is indispensable for their ability to control autoimmune inflammation in the central nervous system. RNAseq analysis uncovered that Lag3-deficient Treg cells upregulate genes highly enriched in metabolic pathways including Myc target genes. We found that Lag3 deficiency resulted in altered Myc expression as well as dysregulated glycolytic activity in Treg cells but not in Th1 effector cells. We also found that Myc expression level directly determines Treg cells’ metabolic profiles as well as their suppressive activity in vivo. Diminishing Myc expression in Lag3-deficient Treg cells reversed glycolytic activity and the impaired suppression, while overexpressing Myc expression in wild type Treg cells was sufficient to disrupt both metabolic activity and in vivo function to alleviate autoimmunity. Collectively, our results uncovered the novel mechanisms by which Lag3 controls Treg cell metabolism and function by regulating Myc expression and metabolic programs.

## Results and Discussion

### Generation of Treg cell-specific Lag3_E1-3_-deficient mice

While there is growing evidence that Lag3 plays a potent immune regulatory function, the precise working mechanism remains largely elusive. Unlike Lag3’s known inhibitory function to downregulate activation of conventional effector T cells, its role in Treg cells is not well understood. To test the role of Lag3 in Foxp3^+^ Treg cell function we generated a mouse model in which the loxp sites are introduced in the *Lag3* gene. Germline Lag3-deficient mice originally developed by Mathis and colleagues targeted the first three exons of the *Lag3* gene that encode the first immunoglobulin-like V type domain containing a proline-rich 30 amino acid extra loop, presumed to be involved in Lag3-MHCII interaction (Huard et al., 1997). As the Lag3-deficient model has been widely used in all the Lag3-focused studies, we chose the same targeting strategy. The exons 1-3 were purposely floxed and subsequently crossed to the Foxp3^Cre^ transgenic mice (Fig 1A). Treg cell-specific exon deletion in the resulting Treg cell-specific Lag3^-/-^ (referred to as Treg^ΔLag3E1-3^ hereafter) was verified by genomic DNA amplification (Fig 1B) as well as by flow cytometry (Fig 1C).

**Fig 1.**
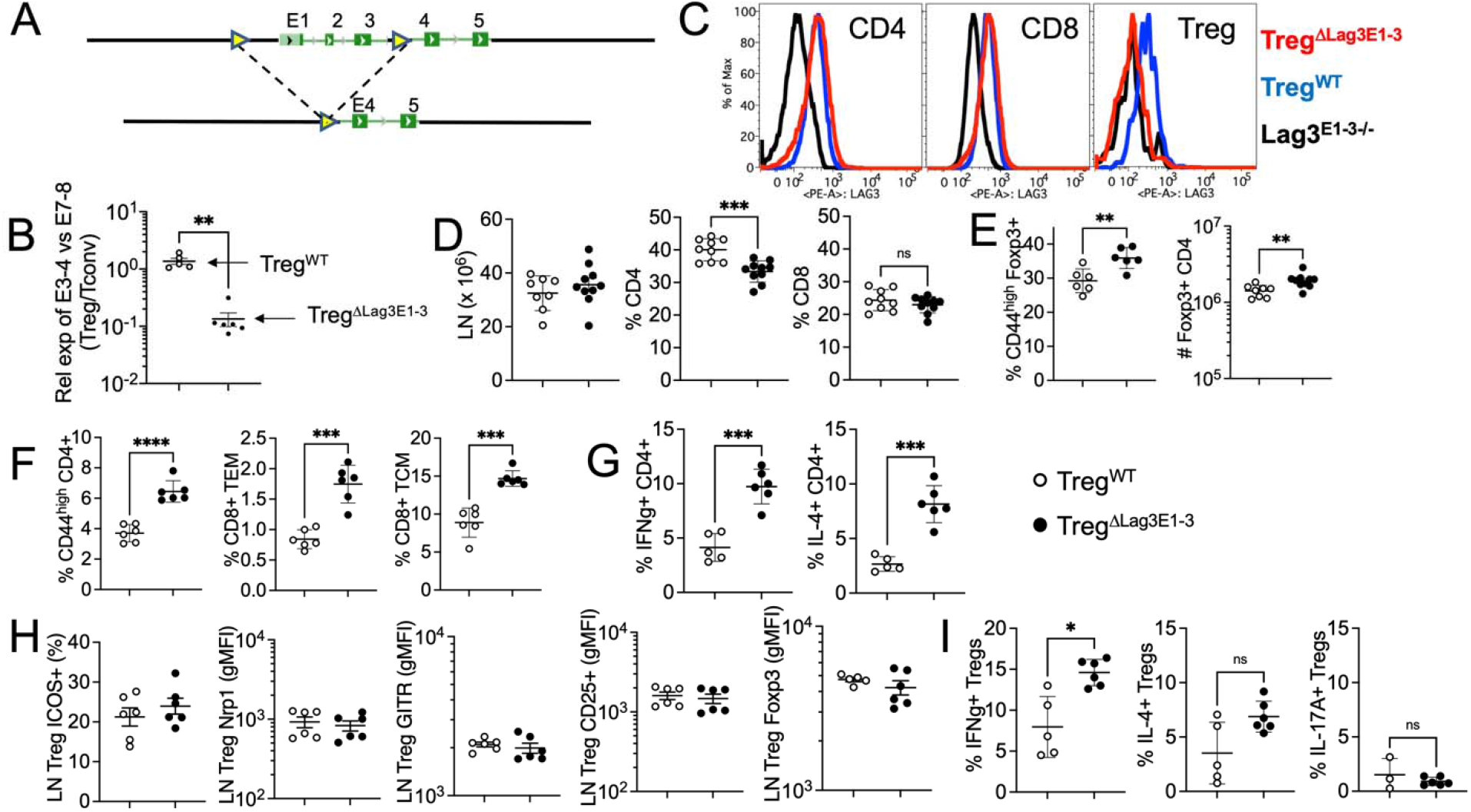
Treg cell-specific Lag3_E1-3_-deficient mice. (**A**) Generation of Treg cell-specific Lag3 exon 1-3 mutant mice by the Cre-loxP system. (**B**) Genomic DNA was extracted from FACS-sorted CD4^+^Foxp3^-^ T cells (Tconv) and CD4^+^Foxp3^+^ T (Treg) cells isolated from Treg^ΔLag3E1-3^ and Treg^WT^ mice. PCR amplifying the exons of the Lag3 gene was performed. Exons 3-4 represent genes targeted in Treg^ΔLag3E1-3^ mice. Amplification of the indicated exon was normalized to the control exon 7-8. (**C**) Lag3 expression of CD4, CD8 and Treg cells in Treg^ΔLag3E1-3^, Treg^WT^ and Lag3^E1-3-/-^ mice. (**D**) Absolute numbers of lymph node (LN) and the proportions of CD4 and CD8 in Treg^ΔLag3E1-3^ and Treg^WT^ mice. (**E**) Absolute numbers of CD44^high^Foxp3^+^ cells and the proportion of CD4^+^Foxp3^+^ T cells in Treg^ΔLag3E1-3^ and Treg^WT^ mice. (**F**) Proportion of CD44^high^CD4^+^, CD8^+^ TEM (CD62L^neg^CD44^high^) and CD8^+^ TCM (CD62L^+^CD44^high^) cells in in Treg^ΔLag3E1-3^ and Treg^WT^ mice. (**G**) Total numbers of IFNγ and IL-4 expressing CD4^+^ T cells in Treg^ΔLag3E1-3^ and Treg^WT^ mice. (**H**) Proportion of ICOS and mean fluorescence intensity (MFI) of Nrp1, GITR, CD25 and Foxp3 on CD4^+^Foxp3^+^ Treg cells in the LN. (**I**) Proportion of IFNγ, IL-4 and IL-17 producing Treg cells. n = 3-6 per group. The results shown are the mean ± SD of individually tested mice from two independent experiments. *p < 0.05, **p < 0.01, ***p < 0.001, ****p < 0.0001 as determined by Mann-Whitney nonparametric test.

Treg^ΔLag3E1-3^ mice were viable and developed normally without gross abnormality. Total cellularity in the secondary lymphoid tissues was comparable to that of wild type control mice (Fig 1D). Interestingly, CD4 T cell proportion was significantly reduced, while CD8 T cell proportion remained unchanged (Fig 1D). We noticed that the proportion as well as total numbers of Treg cells were significantly increased in these mice, suggesting that Lag3 may control Treg cell homeostasis (Fig 1E). Conventional CD4 or CD8 T cells expressing effector/memory phenotypes (CD44^high^) at steady state condition were markedly increased in Treg^ΔLag3E1-3^ mice (Fig 1F). Furthermore, CD4 T cell expression of inflammatory cytokines, such as IFNγ and IL-4, was significantly elevated in Treg^ΔLag3E1-3^ mice (Fig 1G), suggesting the possibility that Lag3-deficient Treg cells may be functionally altered. However, Treg cell phenotypes, determined by ICOS, Nrp1, GITR, CD25, and Foxp3 expression, were comparable regardless of Lag3 expression (Fig 1H). Lag3 deficiency caused Treg cells to express more IFNγ, although IL-4 or IL-17 expression was not affected by Lag3 deficiency (Fig 1I). Therefore, these results suggest that Lag3 deficiency in Treg cells may promote Treg cell expansion at steady state condition and that peripheral T cell homeostasis seems slightly altered possibly due to impaired Treg cell function without overt signs of systemic inflammation.

### Treg cell specific Lag3_E1-3_ deficiency results in greater susceptibility against autoimmune neuroinflammation

To investigate Lag3-dependent Treg cell function during inflammation in vivo, Treg^ΔLag3E1-3^ and Treg^WT^ control mice were induced for EAE. EAE induction was similar between the groups (Fig 2A). However, unlike wild type control mice that partially recover from the initial peak of the disease, Treg^ΔLag3E1-3^ mice continued to display severe disease (Fig 2A). The severity was well reflected by the increased infiltration of CD4 T cells in the CNS tissues (Fig 2B). Analogous to what we have observed from steady state condition (Fig 1E), the proportion as well as total numbers of Treg cells in the CNS were also elevated in Treg^ΔLag3E1-3^ mice (Fig 2B). Therefore, Lag3 may regulate Treg cell expansion and/or survival and Treg cells in Treg^ΔLag3E1-3^ mice may be functionally unable to control inflammatory responses. In fact, Treg cell expression of CD25 and ICOS was significantly downregulated in the mutant Treg cells, although GITR expression remained unchanged (Fig 2C). CNS infiltrating CD4 T cells expressing encephalitogenic cytokines, namely IL-17, IFNγ, and TNFα, as well as IL-10 were substantially elevated in Treg^ΔLag3E1-3^ mice, supporting the severe EAE phenotypes with Treg^ΔLag3E1-3^ mice (Fig 2D). Consistent with the findings, CNS expression of these cytokine genes was markedly increased in Treg^ΔLag3E1-3^ mice (Fig 2E). CNS expression of chemokine genes associated with inflammatory cell infiltration was also dramatically increased (Fig 2F). Similar defects of Lag3_E1-3_-deficient Treg cells to attenuate autoimmune inflammation was also observed in iTreg cells. Adoptive transfer of wild type iTreg cells were fully capable of alleviating severe disease in Foxp3^DTR^ mice with endogenous Treg cell depletion as previously reported (Kim et al., 2019, Fig 2G). On the other hand, Lag3_E1-3_^-/-^ iTreg cells were unable to do so (Fig 2G). Consistent with the disease severity, CNS infiltration of effector CD4 T cells expressing inflammatory cytokines was markedly increased in Lag3_E1-3_^-/-^ iTreg cell recipients (data not shown). Lag3-dependent Treg cell function was further validated in B16 tumor model, as tumor growth was significantly delayed and reduced in Treg^ΔLag3E1-3^ mice (Fig 2H). Taken together, these results strongly suggest that Lag3 plays an important role in Treg cell’s suppressive ability in vivo.

**Fig 2.**
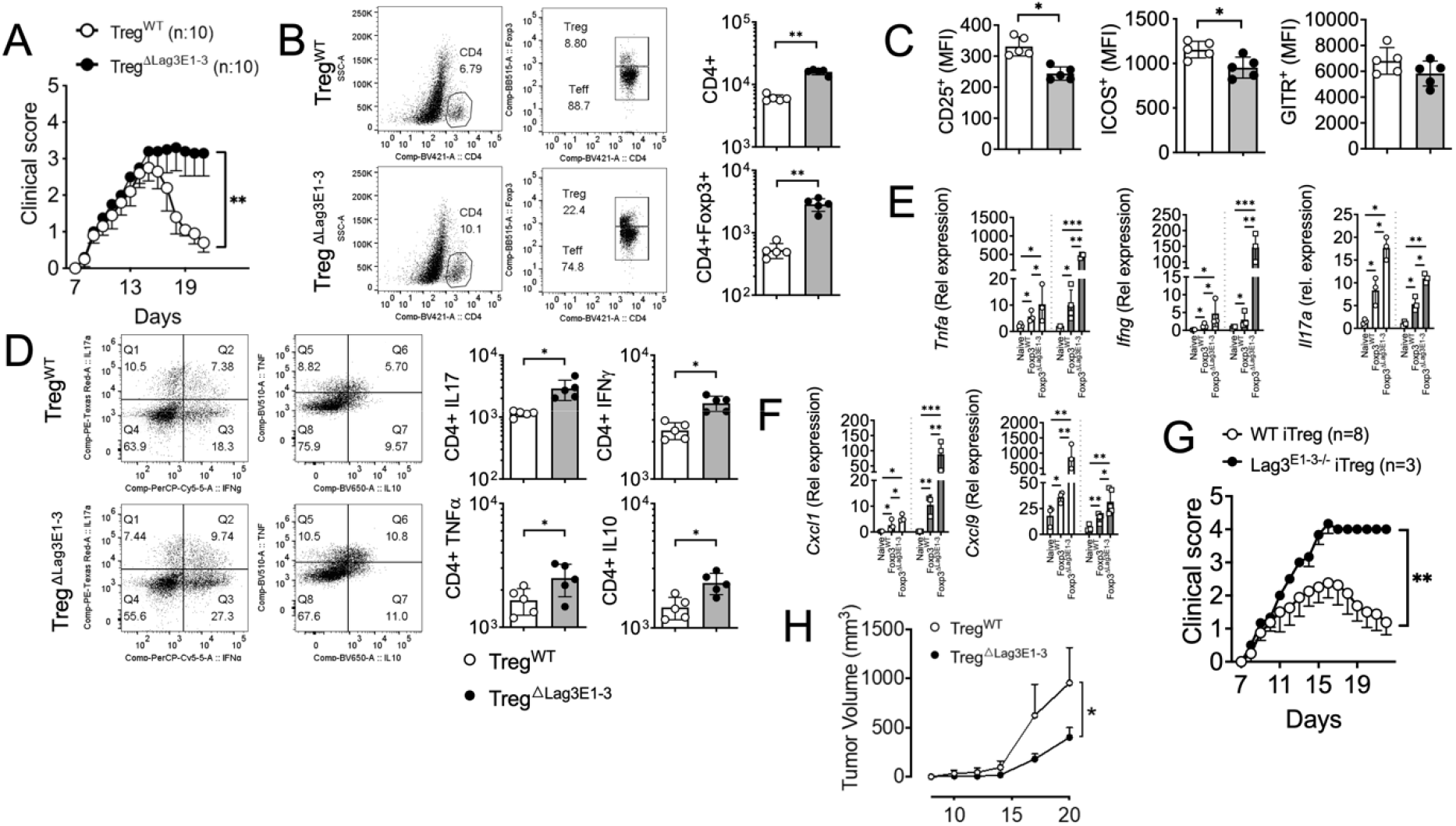
Severe EAE development in Treg cell-specific Lag3_E1-3_-deficient mice. (**A**) Representative clinical scores of Treg^ΔLag3E1-3^ and Treg^WT^ mice. (**B**) Quantification and total number of CNS-infiltrating CD4^+^ and CD4^+^Foxp3^+^ Treg cells at the peak of disease (day 18 postimmunization). (**C**) The mean fluorescence intensity (MFI) of CD25, ICOS and GITR on CD4^+^Foxp3^+^ Treg cells in the CNS at the peak of disease. (**D**) Absolute numbers of GM-CSF, IFNγ, IL-7 and IL-10 expressing CD4^+^ T cells infiltrating the CNS at the peak of disease. (**E-F**) RNA extracted from individual brain and spinal cord of EAE-induced Treg^ΔLag3E1-3^ and Treg^WT^ mice at the peak of the disease were analyzed for the indicated cytokine and chemokine genes by qPCR. (**G**) Foxp3^DTR^ mice were induced with EAE and treated with Diphtheria toxin (DTx) day 7, 8 post induction. 2×10^6^ cell/ml of WT or Lag3^E1-3-/-^ iTreg cells were transferred on day 10 postimmunization. The mice were scored for clinical diseases. (**H**) Quantification of tumor volumes (mm^3^) in Treg^ΔLag3E1-3^ and Treg^WT^ mice following B16 implantation. n = 3-10 per group. The results shown are the mean ± SD of individually tested mice from two independent experiments. *p < 0.05, **p < 0.01, ***p < 0.001, ****p < 0.0001 as determined by Mann-Whitney nonparametric test.

### Lag3_E1-3_-deficiency in Treg cells alters gene expression profiles enriched in metabolic processes

To better understand the mechanisms underlying Lag3-dependent Treg cell functions to control encephalitogenic immunity, we carried out a RNAseq experiment utilizing Treg cells isolated from the CNS at the peak of the disease. Principal component analysis plot demonstrates that Lag3_E1-3_^-/-^ Treg cells displayed distinct clustering compared to wild type Treg cells (Fig 3A). A volcano plot showed that there are ~1000 genes upregulated and >10^4^ genes downregulated in Lag3_E1-3_^-/-^ Treg cells, respectively (Fig 3B). We especially focused on the genes upregulated by Lag3_E1-3_ deficiency through Metascape analysis and found that the top-level Gene Ontology biological processes included those genes enriched in pathways related to metabolic processes (Fig 3C). Geneset enrichment analysis (GSEA) analyses further confirmed that genes upregulated by Lag3_E1-3_ deficiency in Treg cells are highly enriched in cellular metabolic pathways including Myc targets, mTORC1 signaling, oxidative phosphorylation, and fatty acid metabolism (Fig 3D). Other inflammatory pathways including interferon gamma responses, interferon alpha responses, and IL-2-Stat5 signaling were also enriched in Lag3_E1-3_^-/-^ Treg cells. Therefore, these results suggest that Lag3 may play a key role in regulating metabolic profiles as well as cytokine responses in Treg cells.

**Fig 3.**
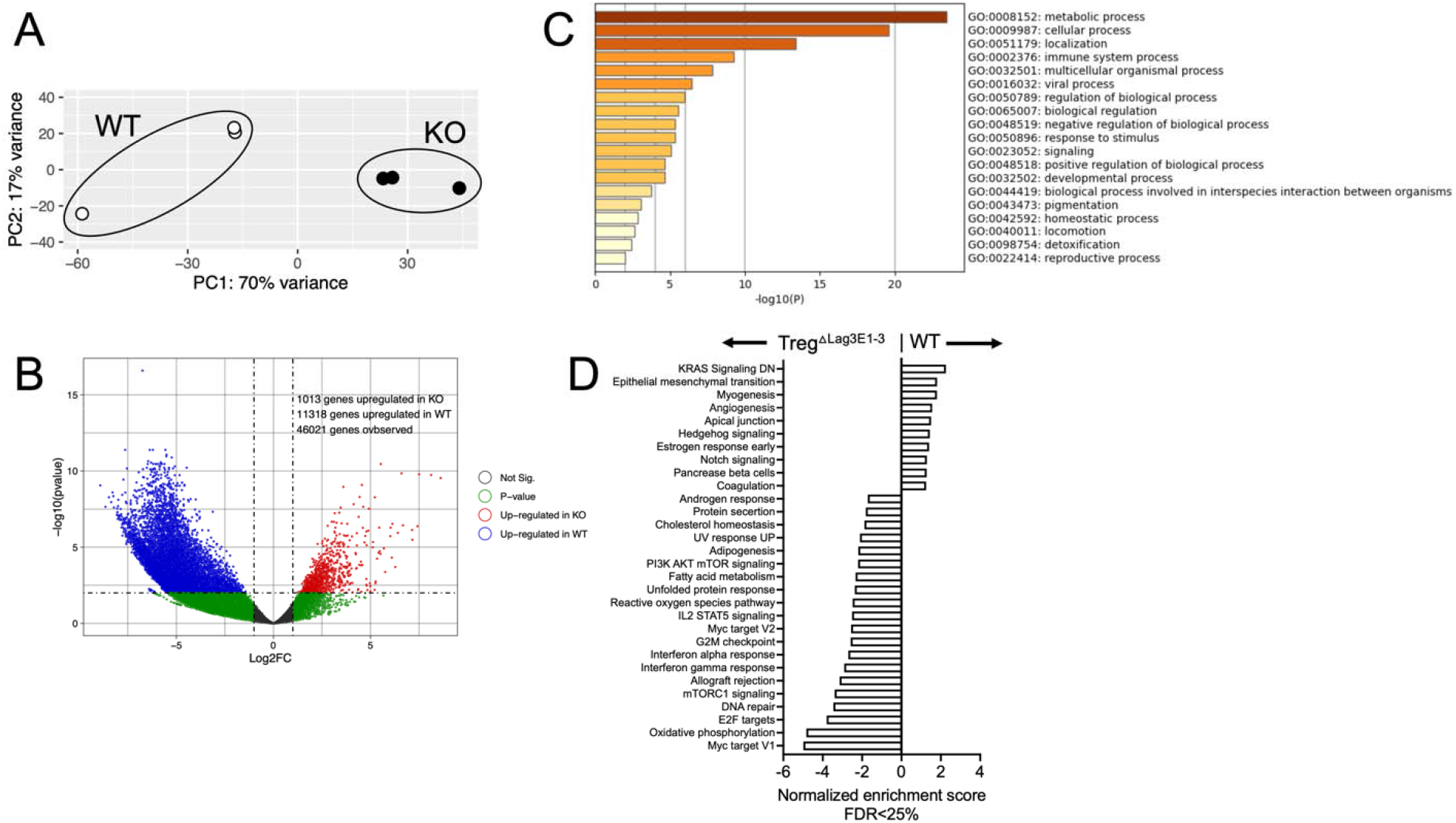
Gene expression profiling of Lag3 ^E1-3^-deficient Treg cells. EAE was induced in Treg^ΔLag3E1-3^ and Treg^WT^ mice by immunization with MOG_35–55_ in CFA as previously described. Treg cells were isolated from the CNS at the peak of the disease, and subjected to RNAseq analysis. (**A**) Principal component analysis of Lag3^E1-3-/-^ and WT Treg cells. (**B**) Volcano plot representing differentially expressed genes between Lag3^E1-3-/-^ and WT Treg cells (n = 3, adjusted p-value threshold = 0.01). (**C**) Gene ontology (GO) analysis biology processes that are associated with genes upregulated in Lag3^E1-3-/-^ Treg cells. (**D**) Enriched pathways in Lag3^E1-3-/-^ and WT Treg cells from Gene Set Enrichment Analysis (GSEA).

### Altered metabolic profiles of Lag3_E1-3_^-/-^ Treg cells

Among the pathways enriched in Lag3_E1-3_^-/-^ Treg cells, the Myc target gene pathways caught our attention (Fig 4A). Myc plays an important role in regulating glycolytic gene expression (Dong et al., 2020). In Treg cells, Foxp3 is known to suppress Myc expression and to downregulate glycolysis in Treg cells (Angelin et al., 2017; Frattini et al., 2012). Because of the dysregulated Myc target gene expression in Lag3_E1-3_^-/-^ Treg cells, we hypothesized that Treg cell metabolism, especially glycolysis, may be altered in the absence of Lag3. To test the possibility, we first measured Myc expression by western blot analysis. Aerobic glycolysis is a metabolic hallmark of activated Th1 type effector cells (Peng et al., 2016). Myc expression of Th1 effector CD4 T cells was evident (Fig 4B). Consistent with the previous report (Angelin et al., 2017), Myc expression of in vitro generated wild type iTreg cells was almost undetectable (Fig 4B). On the other hand, Lag3_E1-3_^-/-^ iTreg cells expressed high level of Myc protein comparable to that of Th1 type effector cells (Fig 4B). Notably, Foxp3 expression of Lag3_E1-3_^-/-^ iTreg cells was comparable to that of wild type Treg cells, raising the possibility that Lag3 but not Foxp3 may be involved in suppressing Myc expression in Treg cells. To examine the consequence of Myc expression in cellular metabolism, we then carried out Seahorse analysis to specifically measure glycolytic activity through extracellular acidification rate (ECAR). As expected, Th1 cells were highly glycolytic compared to Treg cells as demonstrated by elevated ECAR (Fig 4C). Interestingly, Lag3_E1-3_ deficiency did not affect glycolytic activity of Th1 effector cells (Fig 4C). By contrast, Lag3_E1-3_^-/-^ Treg cells displayed markedly increased ECAR compared to that of wild type Treg cells (Fig 4C). Of note, the level of ECAR of Lag3_E1-3_^-/-^ Treg cells was comparable to that of Th1 type effector cells (Fig 4C). Therefore, these results suggest that Lag3 may directly regulate Myc expression and Myc-dependent metabolic activity in Treg cells.

**Fig 4.**
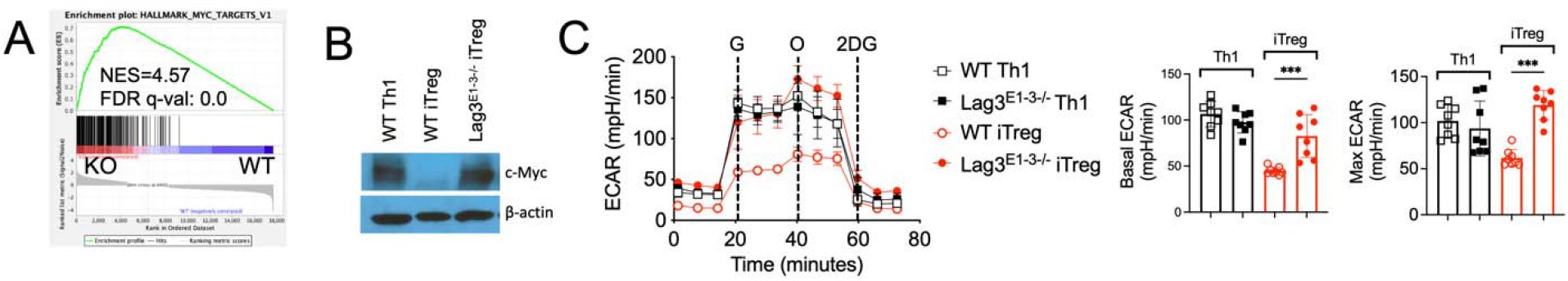
Lag3_E1-3_ deficiency modulates Myc expression in Treg cells. (**A**) GSEA analysis of the Myc target gene set in Treg cells from Treg^ΔLag3E1-3^ and Treg^WT^ mice with EAE. (**B**) Naïve CD4^+^ T cells isolated from Lag3^E1-3-/-^ and control littermates. Cells were activated under Th1 or Treg cell-polarizing conditions and Myc protein expression was determined by western blot analysis. (**C**) Extracellular acidification rate (ECAR) of Lag3^E1-3^ and WT Treg cells or Th1 cells. n = 2-8 per group. The results shown are the mean ± SD of individually tested mice from two independent experiments. ***p < 0.001 as determined by Mann-Whitney nonparametric test.

### Myc expression in Treg cells determines the metabolic activity as well as suppressive functions

Metabolic program of Treg cells is directly wired into their suppressive functions, and increased glycolysis impairs Treg cell suppressive capacity (Gerriets et al., 2016). Since Lag3_E1-3_ deficiency increases Myc target genes and glycolytic activity, we questioned if heightened Myc expression in Lag3_E1-3_^-/-^ Treg cells may be responsible for the defective suppressive function. To test this possibility, Myc expression in Lag3_E1-3_^-/-^ Treg cells was knocked down using a lenti-CRISPR system expressing Cas9 and sgRNA targeting Myc (Sanjana et al., 2014). Myc knockdown was confirmed by western blot analysis (Fig 5A). Importantly, Myc knockdown in Lag3_E1-3_^-/-^ Treg cells reinstated ECAR to the level of WT Treg cells (Fig 5B). To test the impact of Myc knockdown on Treg cell functions, these Treg cells were adoptively transferred into Foxp3^DTR^ mice induced for EAE and treated with DTx to deplete endogenous Treg cells. Myc targeted Lag3_E1-3_^-/-^ Treg cells expressed substantially improved suppressive activity compared to control Lag3_E1-3_^-/-^ Treg cells (Fig 5C). In support, CD4 T cell infiltration in the CNS tissues and accumulation of effector CD4 T cells expressing encephalitogenic cytokines, IFNγ, IL-17, and GM-CSF, were substantially diminished following Myc targeted Lag3_E1-3_^-/-^ iTreg cell transfer (Fig 5D and 5E).

**Fig 5.**
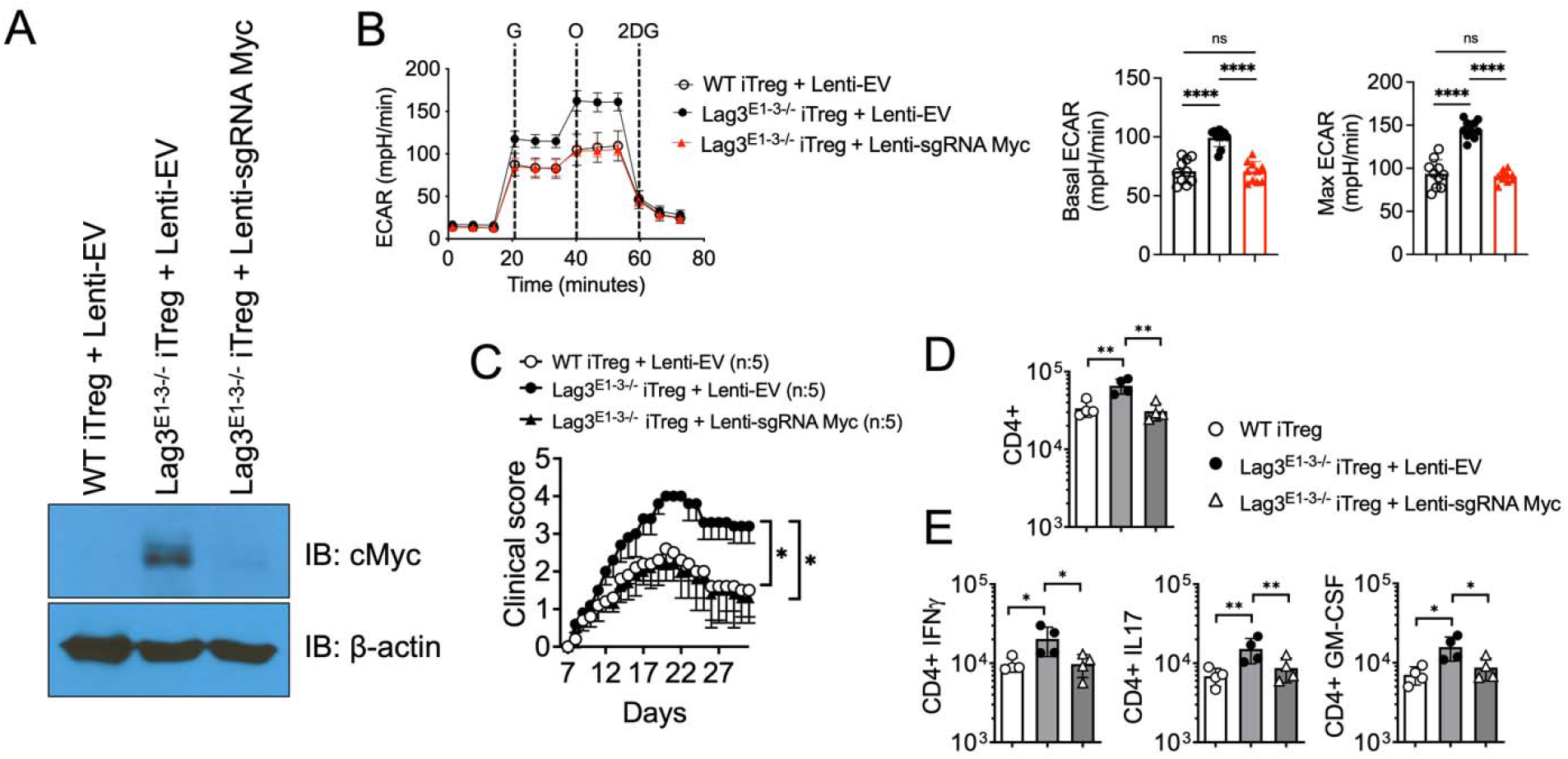
Loss of Myc expression restores metabolic profiles and cellular functions in Lag3^E1-3^-deficient Treg cells. Naïve CD4 T cells from LN of Lag3^E1-3^ and WT mice were transduced with lenti-EV or lenti-sgRNA Myc virus and activated under Treg cell-polarizing for 3 days. (**A**) Myc expression was determined by western blog analysis. (**B**) The ECAR profile, basal and maximum (max) ECAR. (**C**) Foxp3^DTR^ mice were induced for EAE and treated with Diphtheria toxin (DTx). iTreg cells were transferred into the Treg cell-depleted recipients. Disease scores were then measured. (**D-E**) Total number of CNS-infiltrating CD4^+^ T cells and GM-CSF, IFNγ and IL-17 expressing CD4^+^ T cells in the CNS were enumerated at the peak of the disease. The results shown are the mean ± SD of individually tested mice from two independent experiments. n = 2-10 per group. *p < 0.05, **p < 0.01, ****p < 0.0001 as determined by Mann-Whitney nonparametric test.

If Myc knockdown is responsible for the restoration of Treg cell metabolic activity and suppressive function in Lag3_E1-3_^-/-^ Treg cells, it is then possible that enforced Myc expression may disrupt wild type Treg cell metabolism and functions. To test this possibility, we overexpressed Myc in wild type Treg cells and examined its impact. Myc overexpression was validated by western blot analysis (Fig 6A). Myc overexpression in wild type Treg cells significantly increased the ECAR level (Fig 6B). In vivo, Myc overexpressed wild type Treg cells displayed drastically impaired the ability to mitigate autoimmune inflammation (Fig 6C). Consistent with the impaired Treg cell functions, CD4 T cell infiltration and encephalitogenic CD4 T cell accumulation in the CNS was significantly increased in this condition (Fig 6D and 6E). Therefore, these results strongly suggest that Lag3 controls Myc expression and that Myc-associated glycolytic activity plays a key role in mediating Treg cell suppressive function in vivo.

**Fig 6.**
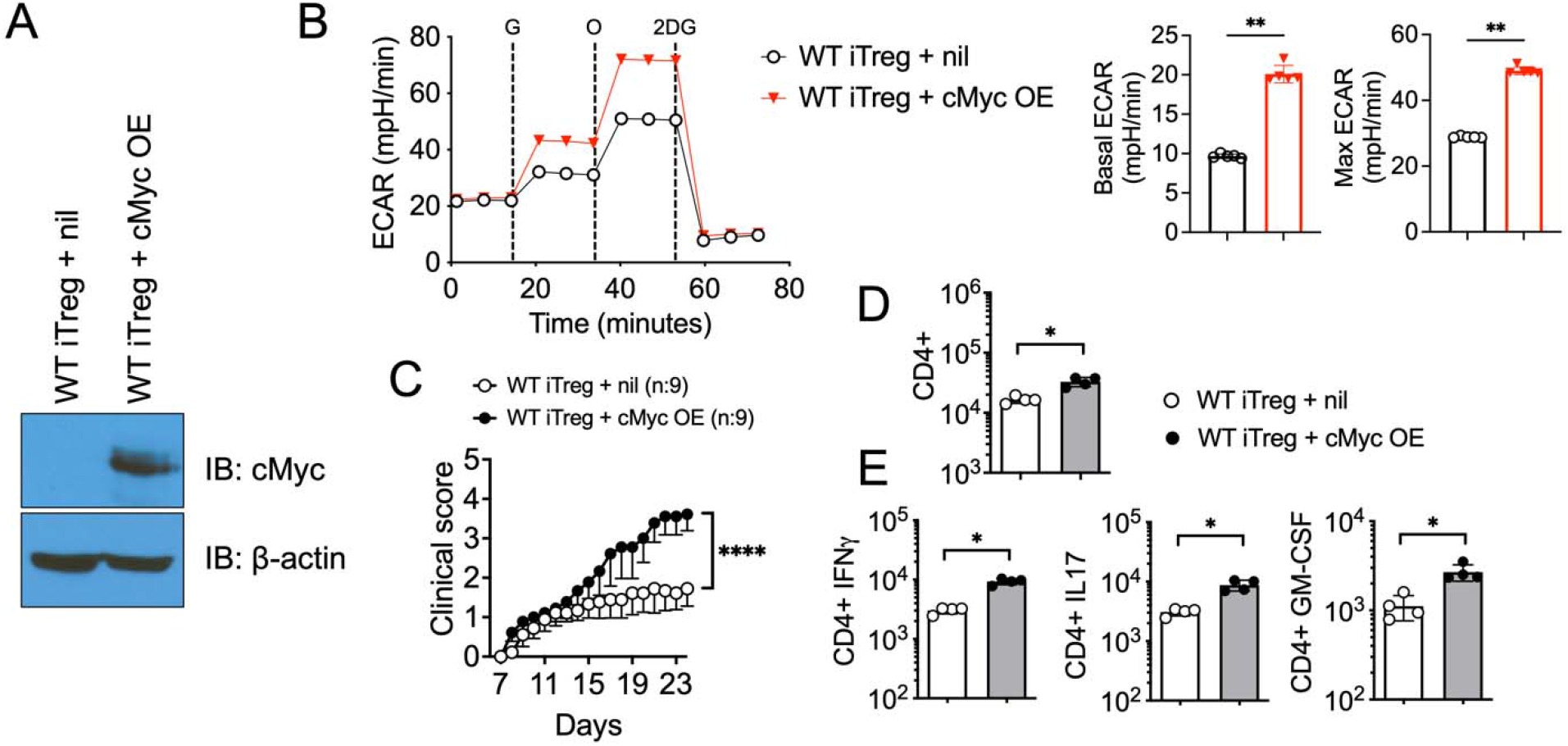
Myc overexpression disrupts the glycolytic activity and function of Treg cells. Naïve CD4 T cells from WT mice were nucleofected with pMXs-cMyc plasmid (Myc overexpression) and activated in vitro under Treg-polarizing conditions. (**A**) Western blot results of Myc expression. (**B**) Measurement of ECAR levels by Seahorse. (**C**) Foxp3^DTR^ mice were induced for EAE and treated with Diphtheria toxin (DTx). iTreg cells were transferred into the Treg cell-depleted recipients. The mice were monitored daily and scored for clinical diseases. (**D-E**) Total number of CNS-infiltrating CD4^+^ T cells and GM-CSF, IFNγ and IL-17 expressing CD4^+^ T cells in the CNS were enumerated at the peak of the disease. The results shown are the mean ± SD of individually tested mice from two independent experiments. n = 2-5 per group. *p < 0.05, **p < 0.01 as determined by Mann-Whitney nonparametric test.

### Concluding remarks

The current study reports several important novel findings. First, our results provide the evidence that Lag3 expression is necessary for adequate Treg cell function to control inflammation in vivo. Lag3 has long been considered critical for Treg cell suppression (Anderson et al., 2016; Gagliani et al., 2013; Huang et al., 2004), although there has been a recent contrasting report that Lag3 instead limits Treg cell function in NOD mice (Zhang et al., 2017). The reason for this discrepancy is not clear. The targeted areas of Lag3 molecule and/or genetic background of the model could play a role. In the present study, we intentionally targeted the same region of the Lag3 gene as the germline Lag3 knockout mice that have widely been used in the previous studies. Given that the targeted first three exons are involved in Lag3-MHCII interaction and in Lag3’s function to inhibit IL-2 production and effector T cell proliferation, our results suggest that the loss-of-function mutation by abrogating Lag3-MHCII interaction seems essential for Treg cells’ ability to control inflammation. Second, our results offer the novel insights of Lag3 in Treg cell metabolism. Treg cell’s differential dependence on cellular metabolism, i.e., glycolysis vs. oxidative phosphorylation, has well been appreciated. Aerobic glycolysis is known to regulate Treg cell proliferation and migration, while it may undermine Treg cells’ suppressive activity (Gerriets et al., 2016). Our result that Lag3_E1-_ 3 deficiency increases the expression of Myc, a key regulator of glycolytic genes, as well as the Myc target genes strongly suggests that Lag3 may limit Myc expression and the subsequent metabolic programs. Myc expression in Treg lineage cells is under tight control by Treg cell specific transcription factor, Foxp3 (Angelin et al., 2017). However, our data suggest that Lag3 may play the major role in regulating Myc expression in Treg cells. The exact signaling pathways connecting Lag3 to Myc remain to be investigated. Third, our result demonstrates that Myc-dependent glycolytic activity directly determines Treg cells’ suppressive activity in vivo. By utilizing lenti-CRISPR or overexpression systems, we demonstrated that Myc-induced aerobic glycolysis directly controls Treg cell function. Collectively, our results offer a novel mechanism by which Lag3 supports Treg cell function by restraining Myc-dependent glycolytic activity.

## Materials and Methods

### Animals

Foxp3^YFPCre^ (B6.129(Cg)-Foxp3tm4(YFP/icre)Ayr/J, #016959) and germline *Lag3^-/-^* mice (B6.129S2-Lag3tm1Doi/J, #026644) were purchased from the Jackson Laboratory (Bar Harbor, ME). The generation of Lag3_E1-3_-floxed mice was commissioned by Biocytogen (Beijing, China). All the mouse protocols were approved by the Institutional Animal Care and Use Committee (IACUC) of the Northwestern University. The mice were maintained under specific pathogen-free conditions at the Northwestern University Feinberg School of Medicine.

### EAE induction

Mice were immunized subcutaneously into the rear flanks with 1:1 emulsion of MOG_35-55_ peptide (300μg, BioSynthesis, Lewisville, TX) in complete Freund’s adjuvant (CFA) containing 5 mg/ml supplemented Mycobacterium tuberculosis H37Ra (Difco, Detroit, MI). Mice were injected with 200 ng pertussis toxin (Sigma, St. Louis, MO) intraperitoneally on day 0 and 2 post immunization. EAE development was monitored and graded on a scale of 0 to 5: 0, no symptoms; 1, flaccid tail; 2, hind limb weakness; 3, hind limb Paralysis; 4, fore limb weakness with hind limb paralysis; and 5, moribund or death.

### Flow Cytometry

Single cell suspensions were prepared from lymph nodes (LN), brain and spinal cord as previously described (Kim et al., 2019). Cells were surface stained with the following Ab: anti-CD4 (RM4–5), anti-CD8 (53-6.7), anti-CD44 (IM7). anti-CD25 (PC61.5), anti-GITR (DTA-1), anti-ICOS (C398.4A), anti-Foxp3 (FJK-16s), anti-Nrp1 (3DS304M). For intracellular cytokine detection, cells were stimulated with PMA (10ng/mL, Millipore-Sigma) and ionomycin (1μM, Millipore-Sigma) for 4 h in the presence of 2μM monensin (Calbiochem, San Diego, CA) during the last 2 h of stimulation. Cells were fixed with 4% paraformaldehyde, permeabilized, and stained with the following Ab: anti-IL-17 (TC11-18H10), anti-IFNγ (XMG1.2), anti-TNFα (TN3-19), anti-IL-10 (JES5-16E3). The Abs were purchased from eBioscience (San Diego, CA), BD Biosciences (San Diego, CA) and BioLegend (San Diego, CA). Stained cells were acquired with a FACSCelesta (BD Biosciences) and analyzed using a FlowJo software (TreeStar, Ashland, OR).

### Seahorse flux analysis

Glycolysis stress test kit and Seahorse XFe96 (Agilent, Santa Clara, CA) were used to evaluate glycolysis. Briefly, cells were glucose-starved in an XF assay medium containing 2 mM glutamine CO^2^-free incubator at 37 °C for 30 min. Extracellular acidification rates (ECAR) were measured by first injecting glucose (1mM) and the cells catabolize glucose into pyruvate via the glycolysis pathway, producing ATP, NADH, water, and protons. Then, oligomycin (2μM) was injected to shift energy production to glycolysis by inhibiting mitochondrial ATP synthesis, and the sharp increase of ECAR after oligomycin injection indicated the level of glycolytic capacity. Finally, 2-deoxyglucosee (2-DG, 2μM), a glucose analog, which inhibited glycolysis through competitive binding to glucose hexokinase, was injected into the wells. Basal ECAR and Max ECAR were calculated according to the manufacturer’s instructions.

### RNAseq

RNA was extracted from Treg cells isolated from the CNS tissues using a RNeasy Micro kit (Qiagen). The quality of all samples was assessed on a Fragment Analyzer electrophoresis system (Agilent). Total RNA was normalized prior to oligo-dT capture and cDNA synthesis with SMART-seq v4 kit (Takara). The resulting cDNA was quantitated using a Qubit 3.0 fluorometer. Libraries were generated using the Nextera XT DNA library prep kit (Illumina). 25 million reads per sample was performed using a HiSeq2500 (Illumina) on a Rapid run flow cell using a 100 base pairs, paired end run. FASTQ files were uploaded and analyzed on Galaxy Project (https://usegalaxy.org). Read qualities were examined by FastQC module (http://www.bioinformatics.babraham.ac.uk/projects/fastqc). RNA sequencing reads were aligned to the genome via STAR version 2.7.8a (https://doi.org/10.1093/bioinformatics/bts635) and the GRCm38.p4(M10) mouse genome annotation file was used as a reference (Dobin et al., 2013). The following parameters were used: pair-ended, mm10 reference genome, 150 bp length of the genomic sequence around annotated junctions, and all other default parameters. Aligned reads were counted using FeatureCount (Liao et al., 2014) with enabled multimapping, multi-overlapping features, and assigned fractions. Counted genes were further analyzed by DESeq2 for differential expression analysis (Love et al., 2014). Metascape analysis (metascape.org) was performed as previously reported (Zhou et al., 2019). GSEA was performed using GSEA software from UC San Diego and Broad Institute (Mootha et al., 2003; Subramanian et al., 2005).

### Western blotting

The protein concentration was adjusted to 30 μg by adding protein lysis buffer (Cell Signaling Technology, Danvers, MA) and 4x loading buffer (Bio-Rad, Hercules, CA). The protein was denatured at 95 °C for 10 min. Proteins were separated by sodium dodecyl sulfate polyacrylamide gel electrophoresis (SDS-PAGE), and transferred to nitrocellulose membrane (Whatman Inc., Florham Park, NJ). After transfer, the membrane was blocked for 1 h. The anti-Myc (Cell Signaling Technology), anti-beta actin (Cell Signaling Technology) primary antibody was added at 4 °C and left overnight. The corresponding secondary antibody (Cell Signaling Technology) was added after washing the membrane on day 2, and was incubated at room temperature for 1 h. The membrane was then developed using an enhanced chemiluminescent HRP substrate (Pierce™ ECL Plus Western Blotting Substrate; Fisher Scientific, Waltham, MA), and visualized after exposure onto HyBlot CL Autoradiography Film (Denville Scientific, Metuchen, NJ) and processing through a SRX-101A film processor (Konica Minolta, Wayne, NJ)

### Quantitative RT-PCR

Total RNA was extracted using a TRIzol reagent according to the manufacturer’s instructions (Invitrogen). RNA concentrations were determined using the NanoDrop (Thermo Scientific). cDNA was synthesized using a MMLV reverse transcriptase (Promega, Madison, WI). Real-time PCR was performed using Radiant qPCR mastermix (Alkali Scientific, Fort Lauderdale, FL) or SYBR Green Master Mix (Applied Biosystems, Waltham, MA) in a QuantStudio 3 Real-Time PCR System (Applied Biosystems). Gene expression was normalized to glyceraldehyde-3-phosphate dehydrogenase *(Gapdh),* before calculating relative gene expression with 2^-ΔCT^ method. The following primers were used:

Taqman primer; *Gapdh* (# 4352661), *Tnfa* (Mm00443258_m1), *Ifng* (Mm01168134_m1) and *Il17a* (Mm00439618_m1).

SYBR primer; *Gapdh* F 5’-ATGCCTGCTTCACCACCTTCT-3’. R 5’-CATGGCCTTCCGTGTTCCTA-3’, *Cxcl1* F 5’-CCGAAGTCATAGCCACACTCAA-3’, R 5’-GCAGTCTGTCTTCTTTCTCCGTTA-3’, *Cxcl9* F 5’-AGAACGGTGCGCTGCAC-3’, R 5’-CCTATGGCCCTGGGTCTCA-3’.

### Myc knockdown and overexpression

For knockdown of Myc using the clustered regularly interspaced short palindromic repeats (CRISPR)/Cas9 technology, sgRNA targeting Myc was cloned into a Cas9-expressing lentiviral transfer vector (lentiCRISPRv2, # 52961, Addgene, Cambridge, MA) following the methods of the Feng Zhang laboratory (Sanjana et al., 2014). To prepare lentiviruses for Myc knockdown, LentiCRISPRv2–sgRNA Myc transfer plasmids was co-transfected with packaging plasmids psPAX2 and pMD2.G (Addgene plasmids #12259 and #12260) into HEK293T cells. Supernatants were harvested and further concentrated using a Lenti-X Concentrator (Takara Bio Inc., Otsu, Japan). Naive CD4^+^ T cells were transduced with lentiviruses and polybrene (8 ug/mL) and activated in the presence of IL-2 (100U/mL) and TGFß (5 ng/mL) to generate inducible Treg cells for 72 h. Myc overexpression was achieved using the Nucleofector™ II/2b (Amaxa, Koelin, Germany) with pMXs-cMyc plasmid (Addgene plasmids #13375). Naive CD4^+^ T cells were resuspended in 100μl nucleofector solution (Amaxa) and add 3μg of pMXs-Myc plasmid into the solution. Then the mixtures were gently transferred to electroporation 0.2 cm cuvettes (Amaxa), placed in the Nucleofector™ II/2b, and nucleofected using the X-001 program. Nucleofected cells were then activated under Treg cell polarizing condition.

### Statistical analysis

Statistical significance was determined by the Mann-Whitney test using a Prism v9.0 software (GraphPad, San Diego, CA). Statistical significance was set at P < 0.05.

